# Integrative x-ray structure and molecular modeling for the rationalization of procaspase-8 inhibitor potency and selectivity

**DOI:** 10.1101/721951

**Authors:** Janice H. Xu, Jerome Eberhardt, Brianna Hill-Payne, Gonzalo E. González-Páez, José Omar Castellón, Benjamin F. Cravatt, Stefano Forli, Dennis W. Wolan, Keriann M. Backus

## Abstract

Caspases are a critical class of proteases involved in regulating programmed cell death and other biological processes. Selective inhibitors of individual caspases, however, are lacking, due in large part to the high structural similarity found in the active sites of these enzymes. We recently discovered a small-molecule inhibitor, 63-*R*, that covalently binds the zymogen, or inactive precursor (pro-form), of caspase-8, but not other caspases, pointing to an untapped potential of procaspases as targets for chemical probes. Realizing this goal would benefit from a structural understanding of how small molecules bind to and inhibit caspase zymogens. There have, however, been very few reported procaspase structures. Here, we employ x-ray crystallography to elucidate a procaspase-8 crystal structure in complex with 63-*R*, which reveals large conformational changes in active-site loops that accommodate the intramolecular cleavage events required for protease activation. Combining these structural insights with molecular modeling and mutagenesis-based biochemical assays, we elucidate key interactions required for 63-*R* inhibition of procaspase-8. Our findings inform the mechanism of caspase activation and its disruption by small molecules, and, more generally, have implications for the development of small molecule inhibitors and/or activators that target alternative (*e.g*., inactive precursor) protein states to ultimately expand the druggable proteome.

## Introduction

Caspases are cysteine proteases responsible for driving many cellular activities and are most well-known for inducing and executing apoptosis. However, caspases are also involved in promoting cellular activation, differentiation, and inflammation events, and their role in these processes is less understood^1–6^. Aberrant proteolysis by this family of proteases can have devastating results, including the proliferation of cancers^7^, neurodegeneration^8,9^, and immunological disorders^10,11^. As such, caspases are heavily regulated within the cellular environment. They are synthesized as a single polypeptide chain with an unformed active site and are maintained in this inactive state until a cellular stimulus triggers proteolysis of the scissile bond after distinct aspartate residues^12,13^.

Efforts to develop selective caspase inhibitors and molecular probes have largely focused on compounds that target the active caspase conformers^14^. Unfortunately, as numerous molecular structures of active caspases have revealed, this family of proteases share highly conserved active site and molecular architecture, leading to significant overlap in substrate specificity^14,15^. This general conservation is supported by *in vitro* studies using short fluorogenic peptide-based substrates and inhibitors with electrophilic warheads^16,17^. Therefore, peptide-based inhibitors, such as the commonly used zVAD-fluoromethyl ketone (zVAD-fmk), are hampered by limited selectivity profiles against both caspase- and non-caspase proteases. Given the rapid rate of activation of most caspases and the subsequent cleavage of downstream executioner caspases, inhibition of active conformers will likely fail to fully block the ensuing consequences of caspase activation. Allosteric inhibitors, such as compounds that target the caspase dimer interfaces have been proposed as an alternative strategy to improve the selectivity profile of caspase inhibitors^18,19^. To date, allosteric caspase inhibitors are only available for caspase-1 and -7.

The promiscuity and incomplete inhibition of active caspase inhibitors could be circumvented by an alternative strategy of targeting procaspases. The maturation of the pro- (inactive or zymogen) enzymes is the primary mechanism of caspase regulation in the cellular environment (Fig. 1A). Although the specific molecular mechanism of activation for individual caspases remains somewhat unresolved, studies have established that,for initiator caspases (*i.e*., caspases-2, -8, -9, and -10), proteolysis is triggered by transient proximity-induced homodimerization followed by intramolecular proteolysis^20,21^. Executioner caspases (*i.e*., caspases-3 and -7) are subsequently subjected to proteolysis by activated initiator caspases. Of the 12 known human caspases, only procaspases-1, -3, and -7 have x-ray crystal structures^22–24^. An NMR structure of the procaspase-8 monomer has also been reported^25^. Consequently, our understanding of the molecular mechanisms of caspase activation, particularly, the determination of whether the processing of caspases occurs *in cis* (intramolecular) or *in trans* (intermolecular) have been limited. Studies have also indicated that the somewhat cryptic enzymatic activity of the unprocessed procaspase likely contributes to a variety of non-apoptotic activities assigned to caspases^25–27^.

**Fig. 1.**
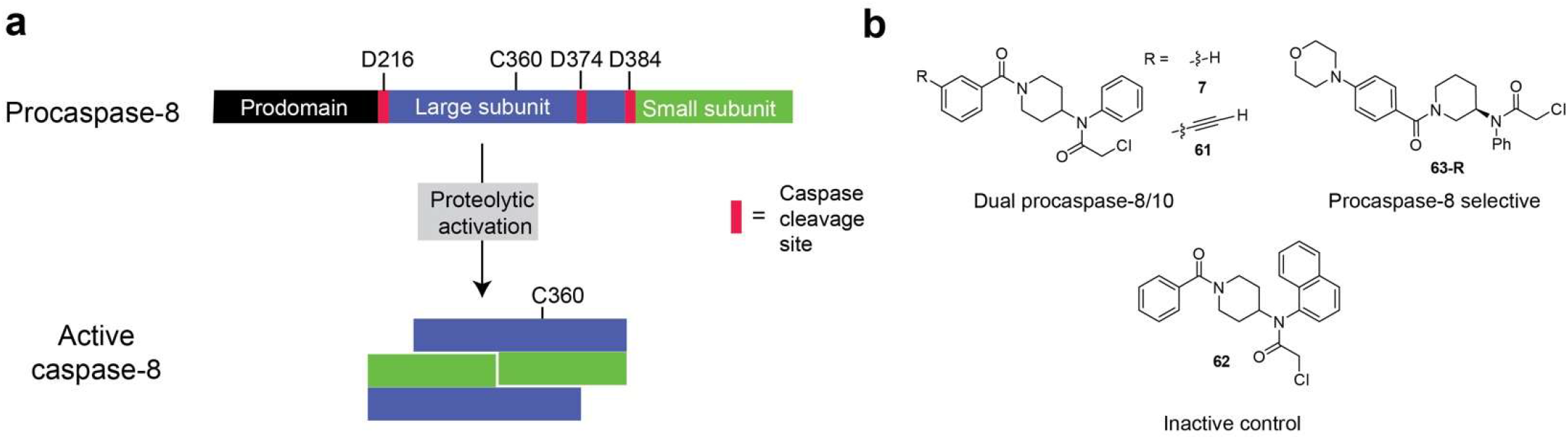
Caspase activation and structures of procaspase inhibitors. **a** General scheme for activation of procaspase-8 by proteolysis after conserved aspartate residues. **b** The structures of caspase-8 lead compounds **7** and **63-*R***, alkyne-containing chemical probe **61** and inactive control compound **62**^28^.

We recently identified several non-peptidic, selective inhibitors of procaspase-8^28^, an initiator caspase that contributes to both extrinsic (Fas ligand-induced) and intrinsic apoptosis^29,30^. These compounds (Fig. 1B) function by irreversible alkylation of the catalytic cysteine in the inactive precursor state of the enzyme, thus blocking activation and subsequent cleavage of downstream executioner caspases-3. The most potent compound, **63-*R***, featured an *alpha*-chloroacetamide electrophile coupled to an *N,N*-disubstituted (*R*)-3-aminopiperidine phenyl core that is further functionalized with a 4-morpholinobenzoyl substituent. Targeting the procaspase conformer is akin to drugging inactive conformations of kinases, a method that has yielded potent and selective inhibitors of several kinases, including the Abl kinase inhibitor imatinib^31–34^.

We report the crystal structure of procaspase-8 in complex with **63-*R***, which reveals that **63-*R*** binds in a pose distinct from that characterized for inhibitors of processed, active forms of caspases. The structure also uncovers large conformational changes in active-site loops that accommodate the intramolecular cleavage events required for caspase-8 processing and activation. To identify and validate key residues involved in ligand recognition and binding, including those not resolved in the crystal structure, we combined molecular modeling with point mutagenesis and binding studies. This hybrid computational-biochemical approach uncovered residues involved in recognition of **63-*R***, another less potent inhibitor **7**, and an alkyne-containing clickable analog of **7** (**61**). Our findings also aided in the rationalization of an inactive, structurally related compound (**62**) (Fig. 1B). We anticipate that the integrated and interdisciplinary strategy described herein will find widespread utility for the identification and functional validation of key structural features missing from crystal structures.

## Results

### X-ray structure of procaspase-8 compared to active caspase-8

We determined the x-ray crystal structure of procaspase-8 (residues 223-479) in complex with **63-*R*** to 2.88 Å resolution (PDB 6PX9) (Fig. 2 and Supplementary Table 1). The final R_cryst_ and R_free_ values were 28.9% and 36.6%, respectively, with 89% of the residues residing the most favored region of the Ramachadran plot (Supplementary Table 1). The structure solution contains 6 molecules per asymmetric unit that form 3 biologically relevant homodimers. Residues 362-388, 409-419, and 453-460 of all 6 subunits lacked interpretable density. All three missing sequences are localized to loops that are exposed to solvent channels, and the missing density suggests these loops are highly flexible.

**Fig. 2.**
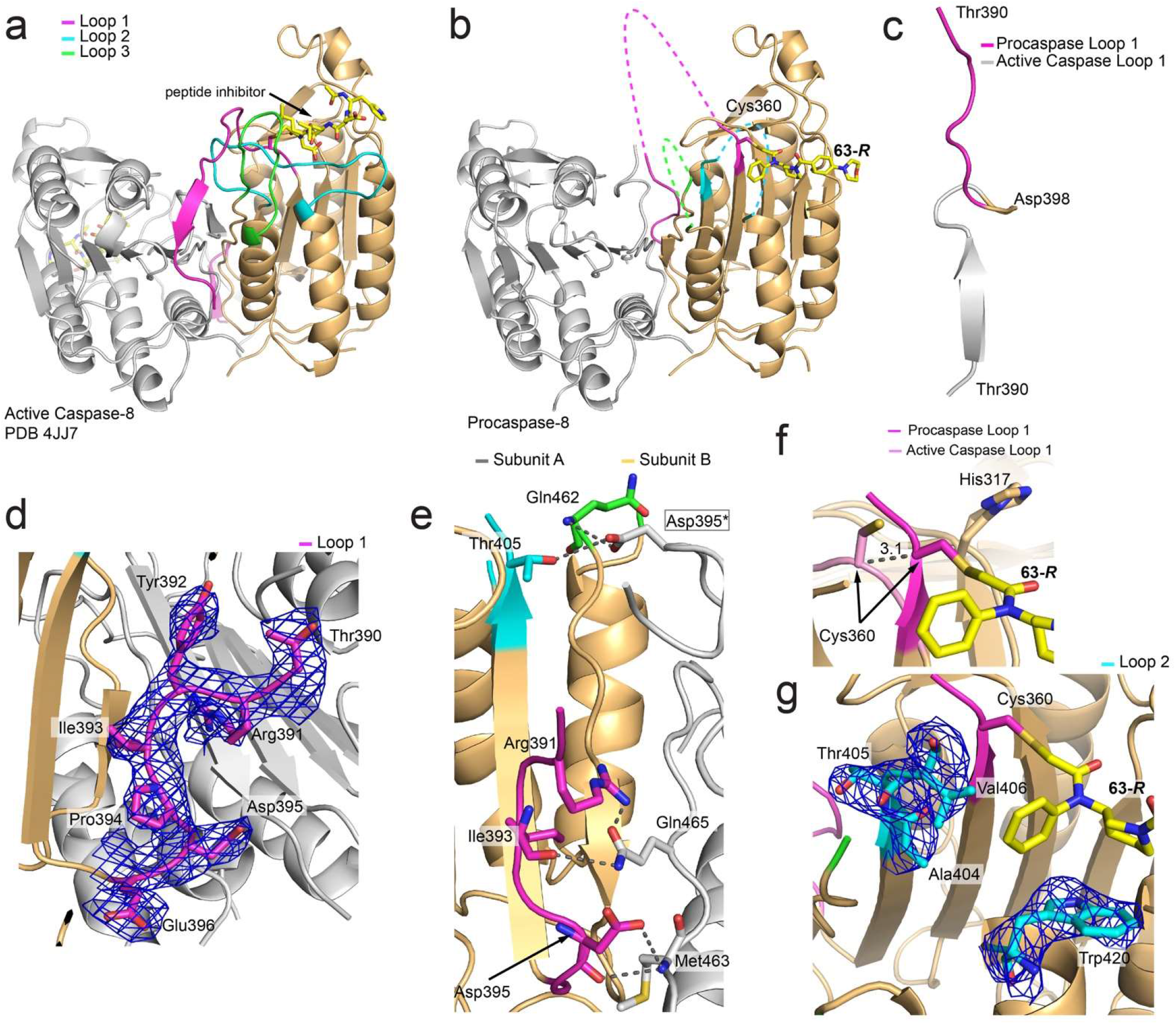
Crystal structure of human procaspase-8. **a** Cartoon representation of homodimeric active caspase-8 bound to covalent inhibitor, Ac-3Pal-D-βhLeu-hLeu-D-AOMK (yellow) shown with the catalytic cysteine (Cys 360) highlighted in magenta and the start and end residues of the three disordered loops, loop 1 (359-396), loop 2, (404-420) and loop 3 (452-462) highlighted in magenta, cyan, and green, respectively, with individual subunits colored tan and grey. **b** The structure of homodimeric procaspase-8 with one chain bound to covalent inhibitor, **63-*R***. Loops, catalytic cysteine, and inhibitor are colored as in ‘**a**’. **c** Overlay of the C-terminal end of loop 1 in active caspase-8 (grey) and procaspase-8 (magenta). Release of loop 1 due to cleavage of the activation linker results in a 180° flip relative to the position of loop 1 in procaspase-8. **d** 2*f*_o_-*f*_c_ density map of well-positioned C-terminal end of loop 1 contoured at 1.0 σ is shown in blue mesh. **e** Potential hydrogen bonds between loops 1, 2, and 3 and the partnering chain at the dimer interface with colors conserved in **a**. **f** Overlay of the catalytic cysteine in active caspase-8 (pink) and procaspase-8 (magenta). **g** 2*f*_o_-*f*_c_ density map of loop 2 contoured at 1.0 σ as in ‘**d**’.

Superposition of procaspase-8 with the structure of active caspase-8 in complex with Ac-3Pal-D-βhLeu-hLeu-D-AOMK (PDB 4JJ7) shows the zymogen core scaffold is highly conserved with the active conformer (Fig. 2A,B). The average main-chain root-mean-square deviation (RMSD) is 0.39 Å (191/256 procaspase-8 Cα). The average RMSD is 0.93 Å for all atoms, with a maximum of 2.43 Å (residues 223-358, 397-403, 421-452, 463-478). The most significant conformational change with respect to the active form is loop 1 (residues 389-396). In the procaspase-8 structure, loop 1 is well positioned over the dimer interface, as demonstrate by clear electron density (Fig. 2C,D), and this orientation is similarly observed for loop 1 in the procaspase-7 crystal structure and procaspase-8 NMR solution structure^22,25^. Upon maturation, loop 1 flips approximately 180° upon cleavage of the activation linker (residues 375-384) and results in the N-terminal region of loop 1 (residues 359-365) contributing key residues to the mature active site, while the C-terminal region of loop 1 (residues 390-396) is solvent exposed (Fig. 2A,C). The primary interactions that position loop 1 are unique to each of the homodimer subunits and thus account for the dimer interface residing on a non-crystallographic symmetry axis. Interestingly, loop 1 of both subunits are positioned by residues within loops 1, 2, and 3 of the opposing subunit. The side chain of Arg391 and the main-chain carbonyl of Ile393 of subunit B are both within hydrogen bonding distance to the side chain of Gln465 of subunit A (Fig. 2E). Likewise, both the side chain and main-chain carbonyl of Asp395 from subunit B interact with the main-chain amide of Met463 from subunit A (Fig. 2E). Conversely, the Asp395 side chain of subunit A forms potential hydrogen bonds with the side chain of Thr405 and main-chain amide from Gln462 of loops 2 and 3, respectively, from subunit B. An important consequence of the intact procaspase-8 loop 1 is the 3.1 Å shift (as measured by Cα) of the catalytic Cys360 residue relative to the activated state (Fig. 2F and Supplementary Fig. 1). The displacement of Cys360 partially occludes the side chain from solvent exposure as well as misaligns the thiol with respect to His317, which is critical to promoting the nucleophilicity of Cys360 in the active caspase-8 structure (Fig. 2F).

In addition to the loop 1 rearrangement, loop 2 (missing residues 409-419) likely has a large conformational shift from the inactive to active conformers. In the active state, residues in loop 2 provide key active site pockets that are required for the recognition and positioning of the non-prime side region of peptide/protein substrates (C-terminal to the scissile bond) (Fig. 2A). Despite much of loop 2 lacking electron density in the procaspase-8 structure (Fig. 2G; see also Supplementary Fig. 1), the N-terminal residues 404-408 of loop 2 form a β-strand that provides a hydrogen-bond network with the neighboring strand containing Cys360 (Fig. 2B and Supplementary Fig. 1). As such, the trajectory of the beginning residues within the loop suggest a distinct conformational orientation compared to the active state. Unfortunately, no residues that comprise loop 3 (residues 453-462) are resolvable in the procaspase structure, suggesting that this region is quite flexible.

### Procaspase-8 interactions with 63-*R*

Crystallization of precursor forms of caspases has likely proven technically challenging due to their structural flexibility, and, in this regard, the procaspase-8 structure appears to have been facilitated by covalent modification with **63-*R***. Electron density was visible for inhibitor **63-*R*** covalently attached to all subunits; however, contiguous density was observed only in subunit B and we opted to model **63-*R*** into this monomer only (Fig. 3A and Supplementary Fig. 2). We also subjected the procaspase-8 crystals to LC-MS/MS analysis to verify that the protease was modified by **63-*R***. Crystals were harvested, washed, solubilized in urea, and digested with trypsin. The trypsin-digested peptides were analyzed by liquid chromatography tandem mass spectrometry using an Q Exactive™ mass spectrometer. The LC-MS/MS analysis confirmed that Cys360 was alkylated by **63-*R*** (Fig. 3 and Supplementary Table 2).

**Fig. 3.**
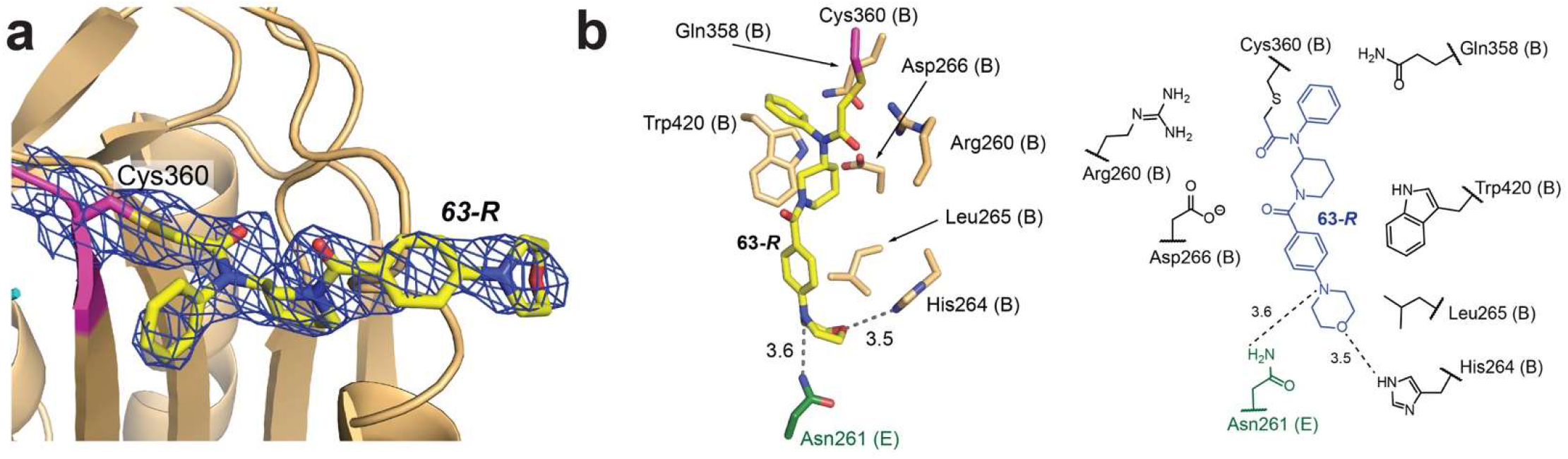
Procaspase-8 in complex with covalent inhibitor **63-*R***. **a** 2*f*_o_-*f*_c_ density map of **63-*R*** bound to catalytic Cys360 contoured at 1.0 σ shown in blue mesh. b Schematic showing the interactions between procaspase-8 and **63-*R*** (yellow carbon in the left stick model and blue in the right schematic). Aside from potential hydrogen bonds between **63-*R*** and His264 as well as a crystal contact from Asn261 of subunit E (green carbon), the rest of the residues form hydrophobic interactions with the inhibitor.

The most significant interaction between **63-*R*** and procaspase-8, aside from the covalent bond between the catalytic Cys360 and the inhibitor, is a weak hydrogen bond formed between the side chain of His264 and the oxygen on the morpholino group (Fig. 3B). A further hydrogen bond is formed with the Asn261 with an adjacent subunit in the crystal lattice (Fig. 3B). Due to the lack of any other polar interactions between the ligand and protein, we hypothesize that the driving force for the procaspase-8-ligand complex is primarily due to hydrophobic interactions. For example, the phenylamine is nestled within a pocket formed by Gln358, Arg260 and Trp420 (Fig. 3B). The piperidine is positioned by the side chain of Trp420 and the benzoyl group has minimal interactions with the protein, leaving the carbonyl exposed to solvent. The morpholino group of **63-*R*** is sandwiched into a pocket formed by residues His264 and Leu265 (Fig. 3B). We believe that the density of the ligand in subunit B is likely continuous due to the additional stability afforded by the potential crystal contact network. The ligand may also reduce the entropy of crystallization by stabilizing the crystal lattice. Superposition of active caspase-8 and procaspase-8 structures show that loop 2 residues 412-418 would directly overlay with the procaspase-8 inhibitor and clearly prevents **63-*R*** from binding the mature caspase.

### Procaspase-8 loop modeling

We next used homology modeling and docking to generate a complete model of the structure of procaspase-8 bound to inhibitors **63-*R***, **7**, as well as the inactive compound **62**. We generated approximately 5,000 models of procaspase-8 to gain insight into the possible conformations of the missing active site loops 1 (residues 362-388), 2 (residues 409-419), and 3 (residues 453-460) and their respective interactions with **63-*R***. The modeling was restricted to the missing regions from the crystal structure and all residues with observable electron density were fixed in position. These models were used to predict **63-*R***-binding residues that lacked electron density in the x-ray structure. Out of the 5,000 models of the homodimer comprised of subunits A and B, we selected the best 1,000 models based on the energy score provided by MODELLER to explain and rationalize the inhibitor SAR and the respective selectivity of the molecules for procaspase-8.

Our analysis confirmed the identities of all residues in close contact (≤ 4 Å) with **63-*R*** and suggested additional contacts from residues within the floppy regions of the loops, including most notably Asn408 (side-chain occupancy: 25.9%; main-chain occupancy: 5.8%); b) Cys409 (21.3%; 6.4%); and c) Asn407 (4.9%; 45.5%) from loop 2 (Fig. 4). These residues are all in close contact with the phenylamide moiety of **63-*R*** (Fig. 4). In addition, we predict a number of loop 2 residues also transiently interact with **63-*R*** with an occupancy ≤ 10 %, such as Pro415, Ser411, Ala416, Arg413 (ranked in decreasing order). We observed that the flexibility of loop 2 likely stems from Gly418, which does not interact directly with **63-*R***. Residues in loops 1 (with the exception of Cys360) and 3 are predicted to have no interactions with **63-*R*** (Fig. 4). It is important to highlight limitations of the models, which are based on the crystallographic template, and therefore only missing loops are considered as flexible, while the rest of the protein and the inhibitor are fixed in position. In non-crystallographic condition, the position of **63-*R*** and the flanking loop 250-266 are expected to present additional flexibility and may limit the accuracy in the binding contributions by specific residues, and the resulting consequences of **63-*R*** binding predicted to occur with mutational analysis. Our prediction accuracy is inversely proportional with respect to their distances from the crystallographically resolved residues. The residue occupancy calculated from the models was used to guide the selection and prioritization of side-chain mutagenesis to rationalize inhibitor SAR via perturbation of procaspase-8 conformational stability/dynamics and/or introduction of steric clashes between **63-*R*** and the procaspase-8 binding site. Both methods of perturbance would result in a measurable and quantitative reduction in ligand binding.

**Fig. 4.**
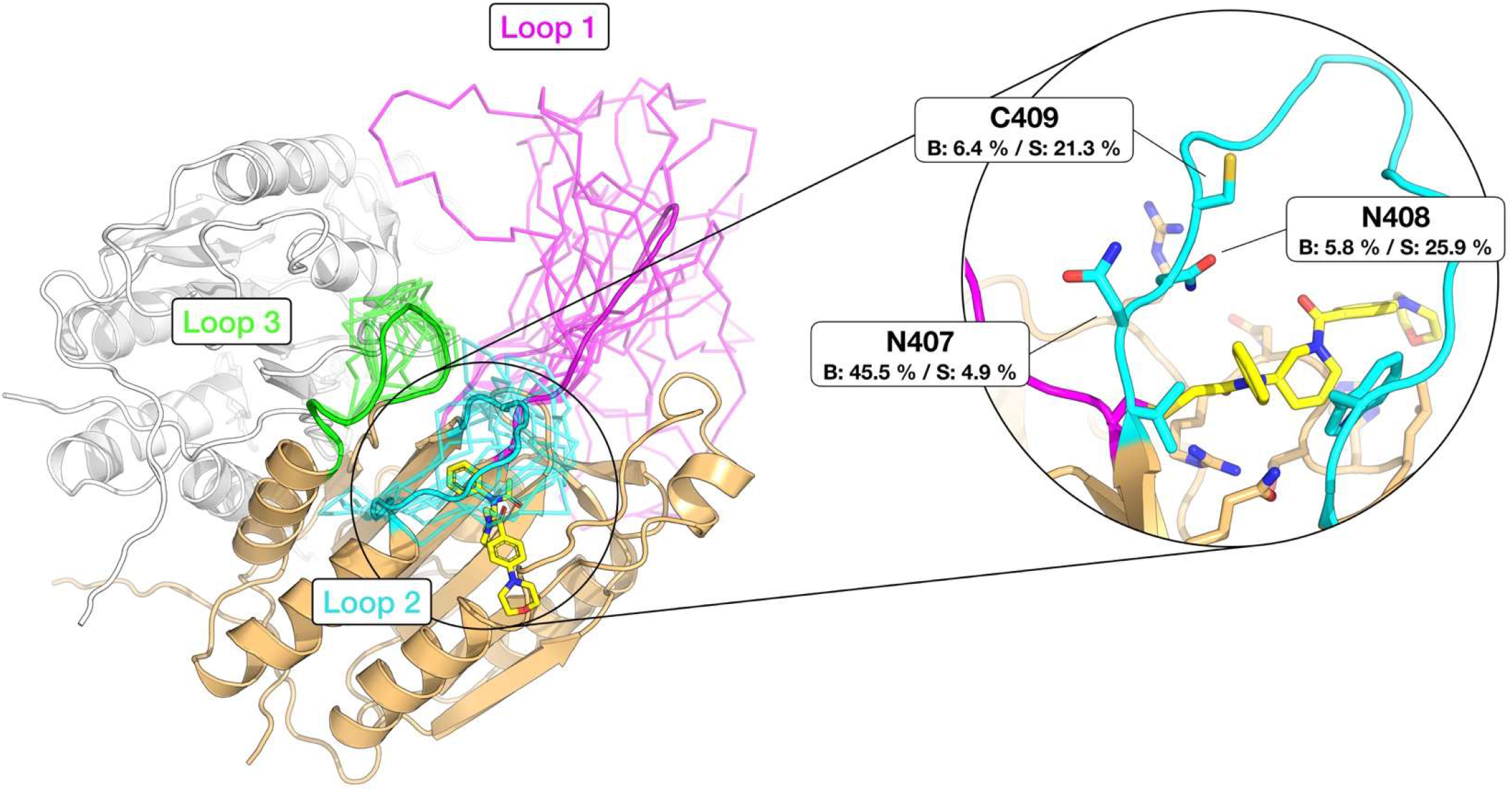
Models of missing loops of procaspase-8 in complex with covalent inhibitor **63-*R***. Ribbon representation of the 10 best models, generated with MODELLER, are shown with the start and end residues of the three disordered loops, loop 1 (359-396), loop 2, (404-420) and loop 3 (452-462) colored in magenta, cyan, and green, respectively, with individual subunits in tan and grey. In cartoon representation and as stick, the representative model of the three disordered loops and residues located around **63-*R***, respectively. The inhibitor **63-*R*** in stick representation with yellow carbon is shown with the backbone and sidechain occupancies of residues N407, N408 and C409 of procaspase-8, as well as the representative model of the loops obtained from the 1000 best models.

### Point mutagenesis and compound binding studies

We next aimed to biochemically validate our structural model (Fig. 4) and determine if the model accurately predicts residues critical for molecular recognition of electrophilic compounds that covalently label Cys360 in the specific active-site conformation formed in procaspase-8 (**63-*R*** and **7**). We also sought to rationalize the previously observed inactivity of compound **62**^28^. We theorized that residues proximal to **63-*R*** would be partially responsible for the potency and selectivity of compound binding and that mutation of these residues would alter compound affinity for procaspase-8. Among the residues within the procaspase-8 active site, the following residues were prioritized for mutational analysis, including: a) Arg260 (predicted to increase the nucleophilicity of Cys360); b) Cys409, the backbone of Asn407, and to a lesser extent Arg258 (predicted to form the compound binding pocket lid); c) Asp266, Gln358 and Trp420 (comprise the bed of the binding site, but do not directly hydrogen bond with either **63-*R*** and **7**) - also notable, Asp266 and Gln358 also contribute to the hydrogen bond network that likely stabilizes the Cys360 thiolate; d) His264 predicted to form a putative H-bond with the morpholino group of the ligand; and Asn261, which was not predicted to interact with the ligand outside of the crystal lattice, and should then serve as a control mutation. All mutant proteins were generated on the uncleavable procaspase-8 construct (see Methods for details). As the Asn407 and Cys409 side chains were hypothesized to form a pocket, both residues were mutated to larger, bulkier groups (N407W and C409W, respectively) to eliminate the pocket and block compound binding. All other resides were mutated to alanine, including R260A, H264A, N261A, D266A, Q358A, and W420A. Unfortunately, the N407W and D266A constructs failed to yield soluble, folded proteins and were removed from further study.

Using a competitive gel-based activity-based protein profiling (ABPP) assay, we assessed the ability of the W420A and the H264A proteins to bind to compounds **7** and **63-*R*** (Supplementary Fig. 3). Briefly, recombinant mutant proteins in cellular lysates were incubated with the indicated compounds at the indicated concentrations for 1h. The mixture was then chased with the alkyne-containing clickable analog of compound **7** (**61** at 10 μM). Direct blockage of procaspase-8 labeling by **61** with pre-incubations in the presence of **63-*R*** or **7** was visualized by Cu(I)-catalyzed azide-alkyne cycloaddition (CuAAC or “click”) conjugation to rhodamine-azide followed by SDS-page and in-gel fluorescence analysis. We were surprised to observe no difference in compound labeling of the W420A compared to the wild-type procaspase-8 (Supplementary Fig. 3A). Next, we chose to investigate the contributions of His264, which can form a labile hydrogen bond with the ligand in the crystal structure. The H264A mutant protein showed comparable compound labeling to that observed for the wild-type procaspase (Supplementary Fig. 3B), confirming the negligible contribution of this interaction to ligand binding.

We hypothesized that the dynamics of the active site loops might contribute to compound binding in a manner that would not be captured by the x-ray structure based on these unexpected results. Given that Gly418 was observed to contribute to loop dynamics in our modeling studies, we postulated that mutation of Gly418 to an alanine would affect the loop dynamics and potentially alter the ligand binding. As such, we added the G418A and G418L mutants to our panel of proteins to test. Competitive gel-based ABPP experiments were performed at a single compound concentration of 10 μM, where **63-*R*** and **7** completely (>90%) or partially (~25%) labeled procaspase-8, respectively, and **62** did not label, consistent with our previous study (Fig. 5A)^28^. As shown previously, **63-*R*** fully competes for labeling of procaspase-8 by **61**, indicating a high-potency labeling event at 10 μM. **7** affords ~75% decrease in labeling by **61**, and **62** exhibits no appreciably competition (Fig. 5B; procaspase-8 gel band). Quite surprisingly given its relative distance from the **63-*R*** (about 9.7 ± 1 Å on average across the models), the G418A mutant exhibited striking changes in SAR across the compounds tested. G418A did not alter labeling by the 4-amino-piperidino containing **7** but did significantly decrease competition of **61** by the 3-amino-piperidino containing **63-*R*** (Fig. 5A-C). While not conclusive, these results indicate that the dynamics and flexibility of loop 2 may contribute to the improved potency of **63-*R*** with respect to **7** (prior calculated IC_50app_ of 0.7 and 5.0 μM, respectively^28^).

**Fig. 5.**
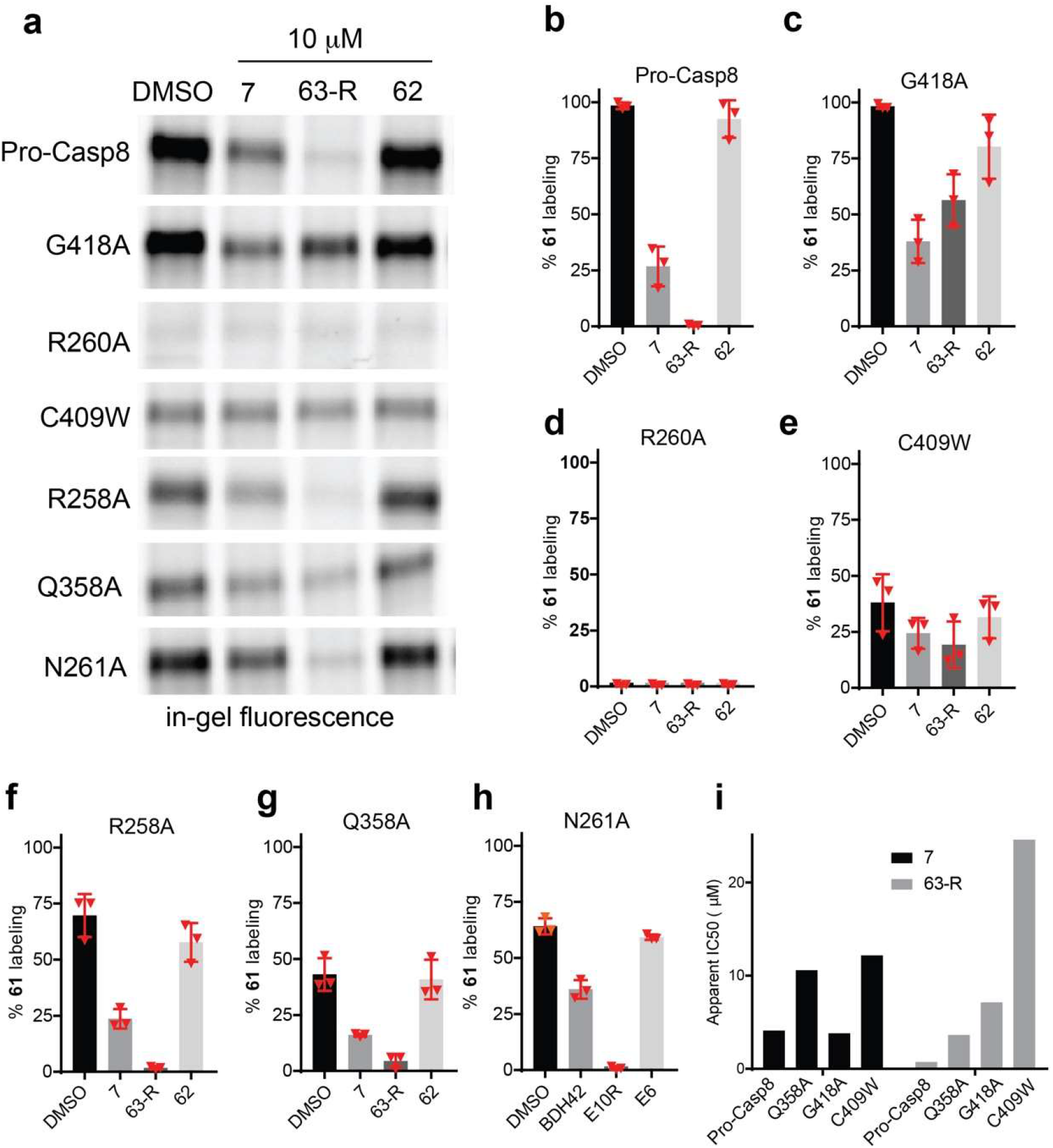
Mutational studies to identify residues that alter probe labeling of procaspase-8. **a** Procaspase-8 recombinant protein (D384A and D394A to prevent activation), which also harbor the indicated mutated residues, were evaluated by gel-based ABPP. Equal concentrations of individual proteins were added to HEK 293T cellular lysates, treated with **7**, **63-*R***, **62**, or vehicle (DMSO) for 1h, followed by labeling with **61** (10 μM) for 1h, “click” conjugation to rhodamine-azide and analysis by SDS-PAGE and in-gel fluorescence. Decrease in fluorescence intensity indicates competition of **61** labeling by compound pre-treatment. **b-h** Quantification of the gel-based data shown in ‘B’. The integrated fluorescence band intensities were quantified and the percentage labeling by **61** was calculated relative to the integrated intensity of DMSO-treated procaspase-8. Shown are the mean and standard deviations derived from three replicate independent experiments for (**b**) procaspase-8, (**c**) G418A, (**d**) R260A, (**e**) C409W, (**f**) R258A, (**g**) Q358A, (**h**) N261A. **I** For the mutant proteins that showed significantly altered compound labeling in ‘**b-c**,’ the apparent half maximal inhibitory concentration (IC_50_) values for compounds **7** and **63-*R*** were calculated for blockade of **61**-labeling from the mean +/- SD of triplicate experiments. Quantification of the decrease in fluorescence compared to vehicle treated samples was calculated from the total integrated intensity of the labeled bands for procaspase-8.

Using our competitive ABPP assay, we next assessed the ability of the procaspase-8 mutant panel (N261A, C409W, Q358A, R260A, and R258A) to bind compounds **7**, **62**, and **63-*R*** (Fig. 5A,D-I and Supplementary Figs. 4 and 5). The R260A mutation, which is soluble and folded comparably to WT, as indicated by CD, showed no appreciable labeling by **7**, consistent with R260 forming a key activating hydrogen bond with the catalytic Cys360 thiol. The bulky C409W mutation, which is predicted to partially occlude the binding site, significantly decreases labeling by **61** and nearly completely blocks competition by pre-treatment with **7** or **63-*R*** and agrees with the loop modeling. Mutation of Arg258, suspected to form part of the lid, did not significantly decrease the apparent potency of compound **7** or **63-*R***. These results are consistent with the low occupancy of Arg258 around **63-*R*** and lack of direct contacts with the ligand. The Q358A mutation, significantly decreased the intensity of protein labeling by **61** and modestly reduced competition of **61** by both **63-*R*** and **7**, which is consistent with our model that Q358, together with other residues, forms the bed beneath the compound, but does not directly hydrogen bond to the ligand. While the N261A mutation caused a slight decrease in **61** labeling, the mutation did not significantly alter the overall binding of **63-*R*** and **7** (Fig. 5H). We confirmed that the purified proteins were folded properly as measured by circular dichroism (CD) (Supplementary Fig. 4A). Similar protein concentrations for all mutant proteins in lysates were further confirmed by immunoblot (Supplementary Fig. 4B).

We next calculated the apparent IC_50_ values of labeling of the Q358A, C409W and G418A mutant proteins by **63-*R*** and **7**. As with the single dose experiments, proteins were spiked into lysates and labeled treated with either **63-*R*** or **7** at a range of concentrations (500 nM – 100 μM) followed by labeling with **61** (10 μM) and the apparent IC_50_ values (Fig. 5I and Supplementary Figs. 6 and 7) were calculated from the competition of labeling by **61**. For the G418A mutation there is no significant change in the IC_50_ of **7**. In contrast, G418A affords a 10-fold increase in the apparent IC_50_ of **63-*R*** from 0.75 μM for procaspase-8 (95% confidence interval (CI), 0.62–0.94) to 7.14 μM for the G418A mutant protein (95% CI, 3.12–14.7), which decreases the potency of **63-*R*** to approximate that of **7**, which labels procaspase-8 with an apparent IC_50_ value of 4.1 μM (95% CI, 2.94-5.80). C409W significantly increases the IC_50_ values of both **7** and **63-*R***, whereas Q358A only modestly alters the apparent IC_50_ values of both **63-*R*** and **7**. Notably, all mutant proteins remained resistant to labeling by control probe **62**, which showed no appreciable competition of labeling by **61**, consistent with our previous studies.

### Pose prediction studies by molecular docking

To further verify and test the x-ray structure and mutagenesis data, we performed silico docking experiments of **63-*R***, **63-*S***, **7**, and **62** with procaspase-8. The covalent dockings were based on the assumption that the four structurally related compounds should all assume similar covalently bound poses. As such, the position of the **63-*R*** phenylamine was assigned as having the highest structural complementarity to the procaspase-8 binding site and was used as a reference anchor for the docking analysis. The predicted position of the inactive **62** 1-naphthyl was used for comparison with the rest of the molecule assuming a similar binding mode as **63-*R*** in the x-ray structure (Fig. 3A). Docking was performed on the 5000 loop models of procaspase-8 generated in absence of **63-*R***, and the best docking score poses were selected for each molecule. Docking success rates were defined as the percentage of correctly placed rings with an RMSD ≤ 1 Å (calculated on the aromatic ring centroids) from the x-ray coordinates. Consistent with our previous biochemical studies, among all docked ligands, the highest success rate was achieved for **63-*R***, which correctly positioned the phenylamine ring in 24% of the 5000 models (Fig. 6A,B). We had previously found that the enantiomer of **63-*R***, compound **63-*S*** was nearly ten-fold less potent^28^. In our docking studies the success rate of **63-*S*** was significantly lower (14%). For **7** and its alkynylated analogue **61**, which in ABPP assays, both show similar potency to **63-*S*** the docking success rate was lower (6 and 8%, respectively). Gratifyingly, the inactive naphthyl compound **62** shows the lowest success rate with 3% (Fig. 6A,C).

**Fig. 6.**
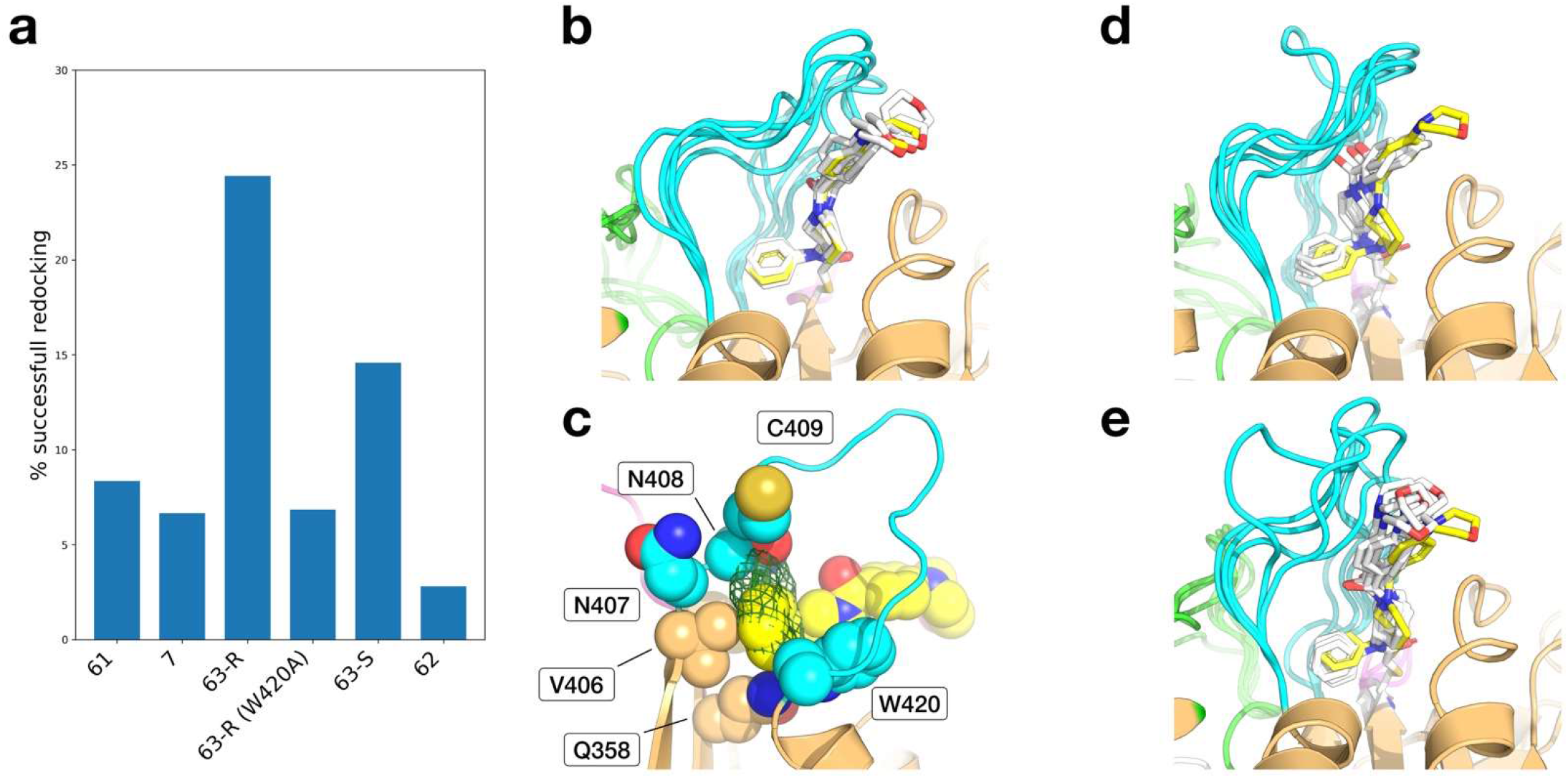
Docking results analysis. **a** Cumulative docking success rates for **61**, **7**, **63-*R***, **63-*S***, and **62** on the 5000 loop models ensemble. **b** The top 5 docking results of **63-*R*** (white carbon) with respect to the experimental coordinates (yellow carbon), with modeled loop 2 conformations in cyan. **c** Docking result of **63-*R*** (yellow spheres) in a representative model, showing how the position of the naphthyl ring in the corresponding position (green mesh) would clash with the residues of the loop 2 and the folded portions of the protein (cyan and tan spheres, respectively). **d,e** The top 5 docking results of compounds **7** and **63-*S***.

The capability of molecules to dock by engaging the hydrophobic pocket of the phenylamine ring appears to partially recapitulate the trend found in the gel-based ABPP assays (Fig. 5). For **62**, modeling shows that only a small fraction of the conformationally accessible loop states are compatible with binding, suggesting that the steric clash of the naphthyl group with the hydrophobic cavity near loop 2 is likely the cause of the observed inactivity in ABPP studies (Fig. 5). Conversely, when modeling the impact of the W420A mutation, we found that the absence of the aromatic side chain has a detrimental effect on docking accuracy, while no effect was found in the mutagenesis experiments (Supplementary Fig. 3). This result indicates that either procaspase-8 undergoes a conformational reorganization to compensate for the W420A mutation, or that the W420A substitution might not impact the initial non-covalent interactions and subsequent covalent alkylation of procaspase-8 by **63-*R***. Also, while experimental data on **61** and **7** show comparable potency with **63-*S***, their docking success rates are smaller than expected (Fig. 6D,E). These shortcomings are likely due to the approximations used in the simulations, including the bias of the loop modeling based on the complex with **63-*R***, the protein maintained as rigid during dockings, and docking scoring function limitations. Overall, the docking results show that loop 2 contributes significantly to ligand binding, and our results underscore the general utility of docking studies in characterizing the potential contributions of disordered regions of proteins to molecular recognition of small molecules.

## Discussion

Our elucidation of the first crystal structure of procaspase-8 in complex with **63-*R*** provides substantial insights into the unique active-site features of precursor caspases that may afford greater specificity for interactions with small-molecule inhibitors compared to the more classical targeting of mature caspase proteases. In the zymogen structure, which is suboptimal for protease activity, key catalytic residues and active site loops are shifted significantly from their positions observed in the structures of the mature, fully activated enzyme, and the locations of these residues overlay closely with the corresponding residues in the structure of procaspase-7 (Fig. 7)^22^. As was observed for the prior procaspase-7 structure, we found significant missing density for three highly flexible active site loops, which we postulate form a lid that partially occludes the active site. One of these loops, loop 1, is cleaved during proteolytic activation of caspase-8. Given the proximity of these loops to the active site and key contributions to caspase activation, we speculated that these loops might, at least in part, be contributing to recognition of **63-*R***. To test this theory, we used molecular modeling to generate a series of high confidence energy minimized models of all three loops in the procaspase-8 conformer. This hybrid structure-modeling approach identified several key residues that were predicted to interact with **63-*R***. Consistent with the observed missing density, our model supported that the loops are highly disordered and can adopt multiple conformations. These results are consistent with prior NMR studies of procaspase-8 that revealed the highly dynamic nature of these loops (Fig. 7). We anticipate that our hybrid method will prove generalizable for a wide range of structures that harbor intrinsically disordered regions.

**Fig. 7.**
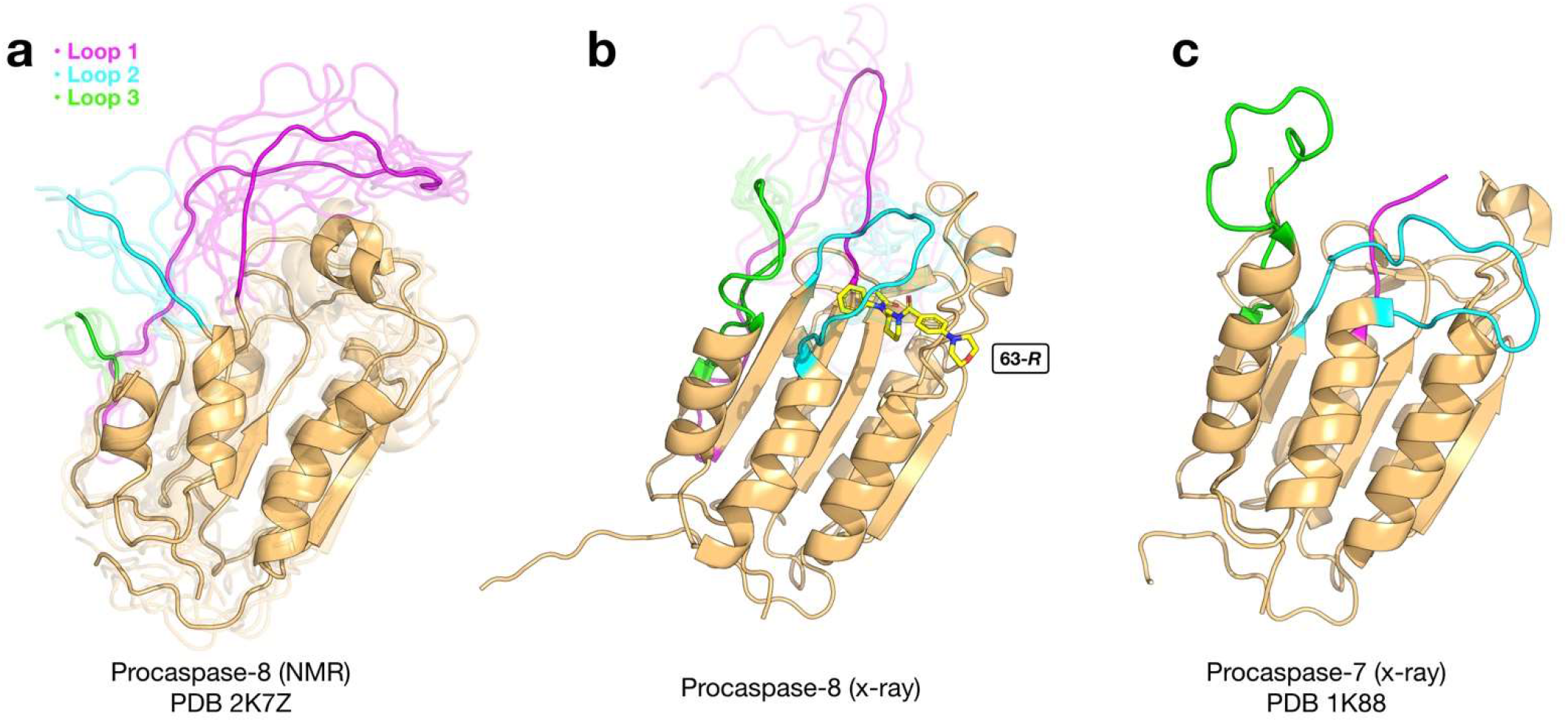
Comparison of different procaspase protein structures. **a** Cartoon representation of the average NMR structure of procaspase-8 (PDB 2k7z), obtained from 20 models, with the first five NMR models represented in transparent. The loops 1, 2, and 3 are colored magenta, cyan and green, respectively. **b** Average structure of procaspase-8 in complex with the inhibitor **63-*R*** (chain B only) in stick representation and colored in yellow, obtained from the 1000 best loop modelling models, with the five best models represented in transparent. **c** X-ray structure of procaspase-7 (PDB 1k88) (chain B only) in cartoon representation with loops 1, 2, and 3 colored magenta, cyan and green, respectively.

A key conclusion from the combined x-ray structure and modeling studies is that nearly all observed interactions between the covalent inhibitor **63-*R*** and the protein are hydrophobic. Only one potential weak hydrogen bond was identified with His264, and mutation of this residue did not alter the potency of **63-*R***, suggesting a negligible role of this interaction in stabilizing ligand binding. This near complete absence of hydrogen bonding interactions is unexpected given the relative potency of the optimized lead inhibitor. Although it is possible that the compound is recognized purely based on hydrostatic interactions, another explanation is that the compound’s pre-alkylation binding pose is distinct from the final pose observed in the x-ray structure. Future structural studies using non-covalent compounds and simulations of the binding process should help to delineate the mechanism of compound binding and subsequent alkylation. However, such studies likely will prove challenging due to the difficulties associated with obtaining diffraction quality crystals for both the pro- and active forms of apo caspase-8, which may be better suited to other structural methods, such as NMR or cryo-electron microscopy.

Our structure-guided mutagenesis study provides a map of critical residues around the procaspase-8 binding site that contribute to inhibitor potency and selectivity. Many of the residues identified by mutagenesis are near to the co-crystallized **63-*R*** and are likely to interact with the compound directly. Quite strikingly, mutation of Gly418, which is located nearly ten angstroms away from the compound, significantly altered the observed SAR, likely due to a reduction in flexibility of loop 2. The relative sensitivity of the protein to mutations at the loop 2 hinge region is, perhaps, not surprising, given the significant repositioning of this loop to form the active conformation. In fact, the portion of loop 2 that forms part of the active site of active caspase-8 and occupies the same space as the covalent inhibitor in the procaspase-8 structure. This suggests that the inhibitor may be recognized by the protein as a mimic of this active site loop. Future studies that use peptide or peptidomimetic molecules that mimic this loop may enable further exploration of this model.

By integrating X-ray structural data with homology modeling, docking, and focused mutagenesis, we developed a multidisciplinary approach to refine and analyze the structure and mode of inhibition of procaspase-8. The resulting structure-model hybrid revealed the large conformational space of disordered active site loops and rationalized the observed SAR of our compound series. Our approach should readily translate to a wide range of other structures and should prove particularly useful for analysis of both reversibly and irreversibly inhibited protein targets. We believe that this structure provides the groundwork for further efforts to elucidate the complete structure of procaspase-8. The SAR and the mutagenesis data presented here lay the foundations for more in-depth studies toward optimized inhibitors with greater activity and proteome-wide specificity. Given the high sequence and structural homology of all caspases, and particularly between caspase-8 and caspase-10, we also anticipate that this structure may reveal key features that distinguish caspase-8 and -10 that can be leveraged to ultimately generate caspase-10 selective inhibitors, which have so far eluded our medicinal chemistry efforts. Such studies will require more extensive biochemical and medicinal chemistry efforts, and it will be essential in the application of more extensive computational approaches (e.g., molecular dynamics simulations and free energy calculations) to better capture the structural heterogeneity of the disordered regions of the protein and characterize their role in ligand recognition.

## Methods

### Procaspase-8 expression and purification

Procaspase-8 is over-expressed as a N-terminal His_6_-tag fusion from *E. coli* BL21DE3pLysS (Strategene) in a pET23b vector. Cells were grown in 2xYT media supplemented with 50 μg/ml ampicillin and chloramphenicol at 37 °C to an OD600 nm of 0.6-0.8. Flasks were then transferred to 12 °C and protein expression was induced with 220 μM IPTG for 16h. Cells were immediately harvested and resuspended in ice cold 100 mM Tris, pH 8.0 and 100 mM NaCl (buffer A) and subjected to 3 cycles of lysis by microfluidization (Microfluidics). The cell lysate was clarified by centrifugation at 14,500 x g for 8 min at 4 °C and soluble fractions were loaded onto a 5 mL HisTrap FF crude Ni-NTA affinity column (GE Amersham) pre-equilibrated with buffer A and eluted with buffer A containing 250 mM imidazole. The eluted protein was immediately diluted 5-fold with buffer B (20 mM Tris, pH 8.0) and purified by anion-exchange chromatography (HiTrap Q HP, GE Amersham) with a 20-column volume gradient to 50% of buffer B containing 1 M NaCl. Fractions corresponding to procaspase-8 were pooled and immediately stored at −80 °C.

### Procaspase-8 mutation

The expression construct that encodes the His_6_-tag zymogen form of caspase-8 (residues 214-479 with D374A D384A, C409S, and C433S mutations) to site directed mutagenesis to generate R260A, G418A, C409W, N261A, R258A mutations. Proteins harboring these mutations were expressed and purified from *E. coli* as described for procaspase-8.

### Western blot analysis

The concentration of recombinant procaspase-8 proteins was calculated by NanoDrop™ and the final protein concentrations adjusted to 500 nM in clarified HEK 293T cellular lysates. The procaspase-containing lysates were then resolved by SDS–PAGE and transferred to PVDF membranes, blocked with 5% milk in TBST and probed with 6x-His Tag Monoclonal Antibody (HIS.H8) (ThermoFisher MA1-21315, 1:3000). Blots were incubated with primary antibodies overnight at 4 °C with rocking and were then washed (3×5min, TBST) and incubated with secondary antibodies (LICOR, IRDye 800LT goat anti-mouse, 1:10,000) for 1h at ambient temperature. Blots were further washed (3× 5min, TBST) and visualized on a BioRad ChemiDoc™ MP Imaging System.

### LC-MS/MS analysis of procaspase-8 63-*R* crystals

Protein crystals were harvested, washed, and solubilized in 50 μL 8M urea (660 mg/mL in PBS). To this was added 10 mM DTT (2.5 μL of 200 mM stock solution) and the reaction was incubated at 65 °C for 15 min following which 20 mM iodoacetamide (2.5 μL of 400 mM stock solution) was added and the reaction incubated for 30 min at 37 °C. The samples were then diluted with 150 μL PBS and to this was added 1 mM CaCl_2_ (2 μL of a 200 mM stock in water) and trypsin (2 μg, Promega, sequencing grade, V5111) and the digestion was allowed to digest overnight at 37 °C with shaking. The samples were then acidified to a final concentration of 5% (v/v) formic acid and desalted using C18 Tips (Pierce 87784), following the manufacturer’s instructions. The samples were analyzed by liquid chromatography tandem mass spectrometry using an Q Exactive™ mass spectrometer (Thermo Scientific) coupled to an Easy-nLC™ 1000 pump. The peptides were eluted on a C18 column with a 5 μm tip (100 μm fused silica, 18 cm) using a 140 min gradient of Buffer B in Buffer A (buffer A: 92% water, 5% acetonitrile, 3% DMSO 0.1% formic acid; buffer B: 5% water, 3% DMSO, 92% acetonitrile, 0.1% formic acid) and a flow rate of 200 nL/min with electrospray ionization of 2.2kV. Data was collected in data-dependent acquisition mode with dynamic exclusion (15 s) and charge exclusion (1,7,8,>8) enabled. Data acquisition consisted of cycles of one full MS scan (400-1800 m/z at a resolution of 70,000) followed by 12 MS2 scans of the nth most abundant ions at resolution of 17,500. The MS2 spectra data were extracted from the raw file using Raw Converter (version 1.1.0.22; available at http://fields.scripps.edu/rawconv/). MS2 spectra were searched using the ProLuCID algorithm (publicly available at http://fields.scripps.edu/yates/wp/?page_id=17) using a reverse concatenated and nonredundant variant of the Human UniProt database (release-2012_11) modified to include the sequence of caspase-8 harboring D374A D384A C409S C433S point mutations. Cysteine residues were searched with a static modification for carboxyamidomethylation (+57.02146 C). Searches also included methionine oxidation as a differential modification (+15.9949 M) and labeling by the compound **63-*R*** (+348.18378 C) as a differential modification^35^.

### Circular dichroism

Circular dichroism (CD) was measured in a JASCO J-715 CD spectrophotometer, scanning 2 times from 250 - 195 nm at 50 nm/min, time constant = 4 sec, bandwidth = 1 nm, slit width = 500 μm. 0.3 mg/mL protein solutions in buffer (25 mM Tris HCl, pH 7.4 and 8.3 mM NaCl) were held in 0.1 cm path length quartz cuvettes. Each secondary structure data set was analyzed via SELCON method against Hennessy and Johnson reference proteins^36,37^.

### Crystallization and x-ray data collection

Inhibitor **63-*R*** was added in a 3-fold molar excess to procaspase-8 (300 uM) in 20 mM Tris, pH 8.0 and 10 mM DTT. The protein:inhibitor mixture was clarified of any precipitant by centrifugation at 3000 x g for 3 min. Crystals were grown by sitting drop-vapor diffusion by mixing equal volumes (1.5 μl) of the procaspase-8:**63-*R*** complex and reservoir solution consisting of 0.08 M imidazole, pH 8.0 and 1 M sodium citrate at 25 °C. Data was collected on a single, flash-cooled crystal at 100 K in a cryoprotectant consisting of mother liquor and 25% glycerol and were processed with HKL2000 in orthorhombic space group P31. The calculated Matthews’ coefficient (V_M_ = 3.14 Å^3^ Da^−1^) suggested six monomers per asymmetric unit with a solvent content of 60%. X-ray data was collected to 2.88 Å resolution on beamline 9.2 at the Stanford Synchrotron Radiation Lightsource (SSRL) (Menlo Park, CA).

### Structure solution and refinement

The procaspase-8 structure was determined by molecular replacement (MR) with Phaser^38^ using the previously published active caspase structure (PDB 4JJ7) as the initial search model. The structure was manually built with WinCoot^39^ and iteratively refined using Phenix^40^ with cycles of conventional positional refinement with isotropic B-factor refinement. Non-crystallographic symmetry (NCS) constraints were applied. The electron density maps clearly identified that **63-*R*** was covalently attached to Cys360 within the active site in subunit B. Water molecules were automatically positioned by Phenix using a 2.5σ cutoff in *f*_o_-*f*_c_ maps and manually inspected. The final R_cryst_ and R_free_ are 28.9% and 36.6%, respectively (Fig. 2 and Supplementary Table 1). The model was analyzed and validated with the PDB Validation Server prior to PDB deposition. Analysis of backbone dihedral angles with the program PROCHECK^41^ indicated that all residues are located in the most favorable and additionally allowed regions in the Ramachandran plot. Coordinates and structure factors have been deposited in the PDB with accession entry 6PX9. Structure refinement statistics are shown in Supplementary Table 1.

### Loop modeling and analysis

Several models of the missing N-terminal residues 217-222, as well as the missing residues in loops 1, 2, and 3, were built using MODELLER 9v21^42^. The homodimeric crystal structure of procaspase-8 subunits A and B within the asymmetric unit with covalently bound **63-*R*** to C360 in chain B was used as the structural template. Due to the lack of density for the side chain of R258 (chain B) and its apparent proximity with the N-terminal region of the flexible loop 2, the side chain orientation was also refined during the loop modeling. Each model was first optimized twice with the variable target function method (VTFM) set to the slow level with 300 steps of conjugate gradients (CG) and an objective function cutoff of 1×10^6^. The models were subsequently refined using molecular dynamics (MD) coupled with simulated annealing (SA), set at the slow level. This protocol was applied to generate 5,000 models of the homodimer consisting of chains A and B.

For the analysis, models were ranked by using the DOPE energy score from MODELLER, and the top 1,000 results were selected for **63-*R*** binding analysis. Using the Python module MDAnalysis^43^, the occupancy of each residue in close contact with the covalently bound **63-*R***, using a distance cutoff of 4 Å, was computed from the selected models. The occupancy was defined as the ratio of the number of models where the residue *i* is in close contact (≤ 4 Å) with the ligand over the total number of selected models. The occupancy values are ranging from 0 % (never in close contact) to 100 % (always in contact). To increase resolution, backbone and sidechain of each residue were considered independently.

### Molecular docking

Using the previously described loop modeling protocol, 5000 additional models were generated in the absence of ligand in order to limit biases toward *63-*R**. In contrast with the loop analysis, all models were used for the docking of **7**, **61**, **62**, **63-*R***, and **63-*S***. The models were prepared for docking by adding hydrogen atoms with REDUCE^44^ at pH 7.0 with Asn, Gln and His sidechains allowed to flip, then following the standard preparation protocol^45^. The 3D coordinates of compound **63-*R*** were taken from the crystallographic structure of procaspase-8, and compounds **7**, **61**, **62**, and **63-*S*** were built with the builder module in PyMOL^46^, using **63-*R*** as reference. The affinity maps were generated using AutoGrid with the standard AutoDock forcefield. The center of the search space was set to position x:-14.0, y:-22.0, z:10), the size of the grid set to 60 x 60 x 60 with a grid spacing of 0.375 Å Å and the smoothing was removed. The ligands were then prepared for covalent docking for AutoDock 4.2.1^47^ on Cys360 following the flexible residue protocol^48^. The standard GA parameters were used to generate 10 docked poses, and the lowest energy pose was selected for each docking.

### Gel-based activity-based protein profiling

25 μL of soluble proteome (1 mg/mL) containing procaspase-8 (500 nM each respectively) was labeled with the indicated concentration of the indicated compounds (1 μL of 25 × stock solution in DMSO) for 1h at ambient temperature followed by labeling with 10 μM of probe 61 (1 μL of 25 × stock solution in DMSO). Subsequently, the samples were subjected to CuAAC conjugation to rhodamine-azide for 1h at ambient temperature. CuAAC was performed with 20 μM rhodamine-azide (50× stock in DMSO), 1 mM tris(2-carboxyethyl) phosphine hydrochloride (TCEP; fresh 50× stock in water, final concentration = 1 mM), ligand (17× stock in DMSO:t-butanol 1:4, final concentration = 100 μM) and 1 mM CuSO4 (50× stock in water, final concentration = 1 mM). Samples were quenched with 10μL 4× SDS-PAGE loading buffer. Quenched reactions were analyzed by SDS-PAGE and visualized by in-gel fluorescence.

### Determination of apparent IC_50_ values

25 μL of proteomes containing the indicated protein at 500 nM final concentration were treated with the indicated compounds for 1h at ambient temperature, labeled with probe **61** for 1h, subjected to CuAAC conjugation to rhodamine-azide, quenched, and analyzed by SDS-PAGE and in-gel fluorescence visualization (n = 3). The percentage of labeling was determined by quantifying the integrated optical intensity of the bands, using ImageJ software14. Nonlinear regression analysis was used to determine the apparent IC_50_ values from a dose-response curve generated using GraphPad Prism 6.

### Statistical analysis

Data are shown as mean ± SD. *P*-values were calculated using unpaired, two-tailed Student’s t-test with values <0.05 considered significant.

## Acknowledgements

This work was supported by the National Institutes of Health R01GM118382 (to DWW) and R01GM069832 (to SF), and the Department of Energy DE-FC02-02ER63421 (to KMB). We thank I. Wilson, R. Stanfield, M. Elsliger, and X. Dai for computational assistance, H. Rosen for access to instrumentation, and the staff of the Stanford Synchrotron Radiation Lightsource.

## Author contributions

KMB, DWW, SF, and BFC conceived of the project. JHX and GEGP solved the x-ray structure. SF and JE conducted computational studies. KMB, BHP and JOC conducted mutagenesis, gel-based ABPP, and CD experiments. KMB, SF, and DW designed experiments and analyzed data. KMB, SF, and DW wrote the manuscript with assistance from JHX and JE.

## References

1. Shalini, S., Dorstyn, L., Dawar, S. & Kumar, S. Old, new and emerging functions of caspases. Cell Death Differ. 22, 526–539 (2014).

2. Li, J. & Yuan, J. Caspases in apoptosis and beyond. Oncogene 27, 6194–6206 (2008).

3. Solier, S., Fontenay, M., Vainchenker, W., Droin, N. & Solary, E. Non-apoptotic functions of caspases in myeloid cell differentiation. Cell Death Differ. 24, 1337–1347 (2017).

4. Fogarty, C. E. & Bergmann, A. Killers creating new life: caspases drive apoptosis-induced proliferation in tissue repair and disease. Cell Death Differ. 24, 1390–1400 (2017).

5. Mukherjee, A. & Williams, D. W. More alive than dead: non-apoptotic roles for caspases in neuronal development, plasticity and disease. Cell Death Differ. 24, 1411–1421 (2017).

6. Nakajima, Y.-I. & Kuranaga, E. Caspase-dependent non-apoptotic processes in development. Cell Death Differ. 24, 1422–1430 (2017).

7. Olsson, M. & Zhivotovsky, B. Caspases and cancer. Cell Death Differ. 18, 1441–1449 (2011).

8. Troy, C. M. & Jean, Y. Y. Caspases: therapeutic targets in neurologic disease. Neurotherapeutics 12, 42–48 (2015).

9. Rehker, J. et al. Caspase-8, association with Alzheimer’s Disease and functional analysis of rare variants. PLoS ONE 12, e0185777 (2017).

10. Su, H. et al. Requirement for caspase-8 in NF-kappaB activation by antigen receptor. Science 307, 1465–1468 (2005).

11. Man, S. M. & Kanneganti, T.-D. Converging roles of caspases in inflammasome activation, cell death and innate immunity. Nat. Rev. Immunol. 16, 7–21 (2016).

12. Cullen, S. P. & Martin, S. J. Caspase activation pathways: some recent progress. Cell Death Differ. 16, 935–938 (2009).

13. Johnson, C. E. & Kornbluth, S. Caspase cleavage is not for everyone. Cell 134, 720–721 (2008).

14. Poreba, M. et al. Small molecule active site directed tools for studying human caspases. Chem. Rev. 115, 12546–12629 (2015).

15. Vickers, C. J., González-Páez, G. E. & Wolan, D. W. Selective detection and inhibition of active caspase-3 in cells with optimized peptides. J. Am. Chem. Soc. 135, 12869–12876 (2013).

16. Julien, O. & Wells, J. A. Caspases and their substrates. Cell Death Differ. 24, 1380–1389 (2017).

17. Thornberry, N. A. et al. A combinatorial approach defines specificities of members of the caspase family and granzyme B. Functional relationships established for key mediators of apoptosis. J. Biol. Chem. 272, 17907–17911 (1997).

18. Hardy, J. A., Lam, J., Nguyen, J. T., O’Brien, T. & Wells, J. A. Discovery of an allosteric site in the caspases. Proc. Nat. Acad. Sci. USA 101, 12461–12466 (2004).

19. Scheer, J. M., Romanowski, M. J. & Wells, J. A. A common allosteric site and mechanism in caspases. Proc. Nat. Acad. Sci. USA 103, 7595–7600 (2006).

20. Boatright, K. M. et al. A unified model for apical caspase activation. Mol. Cell 11, 529–541 (2003).

21. Boatright, K. M. & Salvesen, G. S. Mechanisms of caspase activation. Curr Opin Cell Biol 15, 725–731 (2003).

22. Chai, J. et al. Crystal structure of a procaspase-7 zymogen: mechanisms of activation and substrate binding. Cell 107, 399–407 (2001).

23. Thomsen, N. D., Koerber, J. T. & Wells, J. A. Structural snapshots reveal distinct mechanisms of procaspase-3 and -7 activation. Proc. Nat. Acad. Sci. USA 110, 8477–8482 (2013).

24. Elliott, J. M., Rougé, L., Wiesmann, C. & Scheer, J. M. Crystal Structure of Procaspase-1 Zymogen Domain Reveals Insight into Inflammatory Caspase Autoactivation. J. Biol. Chem. 284, 6546–6553 (2009).

25. Keller, N., Mares, J., Zerbe, O. & Grutter, M. G. Structural and biochemical studies on procaspase-8: new insights on initiator caspase activation. Structure 17, 438–448 (2009).

26. Pop, C., Fitzgerald, P., Green, D. R. & Salvesen, G. S. Role of proteolysis in caspase-8 activation and stabilization. Biochemistry 46, 4398–4407 (2007).

27. Philip, N. H. et al. Activity of Uncleaved Caspase-8 Controls Anti-bacterial Immune Defense and TLR-Induced Cytokine Production Independent of Cell Death. PLoS Pathog 12, e1005910 (2016).

28. Backus, K. M. et al. Proteome-wide covalent ligand discovery in native biological systems. Nature 534, 570–574 (2016).

29. Medema, J. P. et al. FLICE is activated by association with the CD95 death-inducing signaling complex (DISC). EMBO J 16, 2794–2804 (1997).

30. Engels, I. H. et al. Caspase-8/FLICE functions as an executioner caspase in anticancer drug-induced apoptosis. Oncogene 19, 4563–4573 (2000).

31. Nagar, B. et al. Crystal structures of the kinase domain of c-Abl in complex with the small molecule inhibitors PD173955 and imatinib (STI-571). Cancer Res. 62, 4236–4243 (2002).

32. Capdeville, R., Buchdunger, E., Zimmermann, J. & Matter, A. Glivec (STI571, imatinib), a rationally developed, targeted anticancer drug. Nat. Rev. Drug Discov. 1, 493–502 (2002).

33. van Montfort, R. L. M. & Workman, P. Structure-based design of molecular cancer therapeutics. Trends Biotechnol 27, 315–328 (2009).

34. Lin, Y.-L., Meng, Y., Jiang, W. & Roux, B. Explaining why Gleevec is a specific and potent inhibitor of Abl kinase. Proc. Nat. Acad. Sci. USA – (2013). doi:10.1073/pnas.1214330110

35. Tabb, D. L., McDonald, W. H. & Yates, J. R. DTASelect and Contrast: tools for assembling and comparing protein identifications from shotgun proteomics. J. Proteome Res. 1, 21–26 (2002).

36. Sreerama, N. & Woody, R. W. A self-consistent method for the analysis of protein secondary structure from circular dichroism. Anal. Biochem. 209, 32–44 (1993).

37. Hennessey, J. P. & Johnson, W. C. Information content in the circular dichroism of proteins. Biochemistry 20, 1085–1094 (1981).

38. McCoy, A. J. et al. Phaser crystallographic software. J. Appl. Crystallogr. 40, 658–674 (2007).

39. Emsley, P., Lohkamp, B., Scott, W. G. & Cowtan, K. Features and development of Coot. Acta Crystallogr. D Biol. Crystallogr. 66, 486–501 (2010).

40. Adams, P. D. et al. PHENIX: a comprehensive Python-based system for macromolecular structure solution. Acta Crystallogr. D Biol. Crystallogr. 66, 213–221 (2010).

41. Laskowski, R. A., MacArthur, M. W., Moss, D. S. & Thornton, J. M. PROCHECK: a program to check the stereochemical quality of protein structures. J. Appl. Crystallogr. 26, 283–291 (1993).

42. Webb, B. & Sali, A. in Protein Structure Prediction 1137, 1–15 (Humana Press, New York, NY, 2014).

43. Michaud-Agrawal, N., Denning, E. J., Woolf, T. B. & Beckstein, O. MDAnalysis: a toolkit for the analysis of molecular dynamics simulations. J. Comput. Chem. 32, 2319–2327 (2011).

44. Word, J. M., Lovell, S. C., Richardson, J. S. & Richardson, D. C. Asparagine and glutamine: using hydrogen atom contacts in the choice of side-chain amide orientation. J. Mol. Biol. 285, 1735–1747 (1999).

45. Forli, S. et al. Computational protein-ligand docking and virtual drug screening with the AutoDock suite. Nat. Protoc. 11, 905–919 (2016).

46. Lill, M. A. & Danielson, M. L. Computer-aided drug design platform using PyMOL. J. Comput. Aided Mol. Des. 25, 13–19 (2011).

47. Morris, G. M. et al. AutoDock4 and AutoDockTools4: Automated docking with selective receptor flexibility. J. Comput. Chem. 30, 2785–2791 (2009).

48. Bianco, G., Forli, S., Goodsell, D. S. & Olson, A. J. Covalent docking using autodock: Two-point attractor and flexible side chain methods. Protein Sci. 25, 295–301 (2016).

