## Supplementary Information for "Integrative x-ray structure and molecular modeling for the rationalization of procaspase-8 inhibitor potency and selectivity"

**Table of Contents**

**S2** **Supplementary Table 1**. X-ray data collection and structure refinement statistics of procaspase-8 in complex with **63-*R***.

**S3** **Supplementary Fig. 1**. Overlay of pro- and active caspase-8 loops 1 and 2.

**S4** **Supplementary Fig. 2**. Simulated-annealing omit map density for **63-*R***.

**S5** **Supplementary Table 2.** Modification of procaspase-8 crystals by **63-*R***.

**S6** **Supplementary Fig. 3.** Competitive ABPP gels of the W420A and H264A mutated forms of procaspase-8.

**S7** **Supplementary Fig. 4.** Circular dichroism and immunoblot data of procaspase-8 mutants.

**S8** **Supplementary Fig. 5.** Representative full-length gels for competitive labeling experiments.

**S8** **Supplementary Fig. 6.** Calculated apparent $IC_{50}$ for labeling of procaspase-8 mutations by **63-*R*** and **7**.

**S10** **Supplementary Fig. 7.** Representative full-length gels for $IC_{50}$ competitive labeling experiments.

**S11** **Supplementary References**

**Supplementary Table 1.** X-ray data collection and structure refinement statistics of procaspase-8 in complex with ***63-R***.

| **Structure** | 6PX9 |
|---|---|
| Space group | P $3_1$ |
| Cell dimensions | |
| a, b, c; Å | 101.3, 101.3, 175.5 |
| α, β, γ; ° | 90, 90, 120 |
| Data Processing | |
| Resolution, Å (outer shell) | 50.0-2.88 (2.93-2.88) |
| Completeness, % | 99.5 (99.5) |
| Unique reflections | 45,476 (2,243) |
| Redundancy | 4.5 (4.4) |
| $R_{meas}$ (%)[a] | 33.6 (150) |
| $R_{merge}$ (%)[b] | 29.7 (132) |
| $R_{p.i.m.}$ (%)[c] | 15.6 (70.5) |
| Average I / Average σ (I) | 8.1 (1.7) |
| $CC_{1/2}$ | 67.9 (17.4) |
| **Refinement** | |
| Resolution, Å (outer shell) | 50.0-2.88 (2.94-2.88) |
| No. reflections (test set)[d] | 45,343 (2,253) |
| $R_{cryst}$ (%)[e] | 28.9 (44.2) |
| $R_{free}$ (%) | 36.6 (49.4) |
| Protein atoms / Waters | 9,845 / 4 / 30 |
| CV coordinate error (Å)[f] | 0.90 |
| RMSD bonds (Å) / angles ° | 0.003 / 0.677 |
| B-values protein/waters/ligands (Å$^2$) | 44 / 39 / 43 |
| Ramachandran Statistics (%) | |
| Preferred | 89.2 |
| Allowed | 10.8 |
| Outliers | 0 |

[a]$R_{meas} = \{\Sigma_{hkl}[N/(N-1)]1/2\Sigma_i|I_{i(hkl)} - \langle I_{(hkl)}\rangle|\} / \Sigma_{hkl}\Sigma_i I_{i(hkl)}$, where $I_{i(hkl)}$ are the observed intensities, $\langle I_{(hkl)}\rangle$ are the average intensities and N is the multiplicity of reflection hkl. [b]$R_{merge} = \Sigma_{hkl}\Sigma_i|I_{i(hkl)} - \langle I_{(hkl)}\rangle| / \Sigma_{hkl}\Sigma_i I_{i(hkl)}$ where $I_{i(hkl)}$ is the i$^{th}$ measurement of reflection h and $\langle I_{(hkl)}\rangle$ is the average measurement value. [c]$R_{p.i.m.}$ (precision-indicating $R_{merge}$) = $\Sigma_{hkl}[1/(N_{hkl} - 1)]^{1/2}\Sigma_i|I_{i(hkl)} - \langle I_{(hkl)}\rangle| / \Sigma_{hkl}\Sigma_i I_{i(hkl)}$. [d]Reflections with I > 0 were used for refinement[1-3]. [e]$R_{cryst} = \Sigma_h||F_{obs}|-|F_{calc}||/\Sigma|F_{obs}|$, where $F_{obs}$ and $F_{calc}$ are the calculated and observed structure factor amplitudes, respectively. $R_{free}$ is $R_{cryst}$ with 5.0% test set structure factors. [f]Cross-validated (CV) Luzzati coordinate errors.

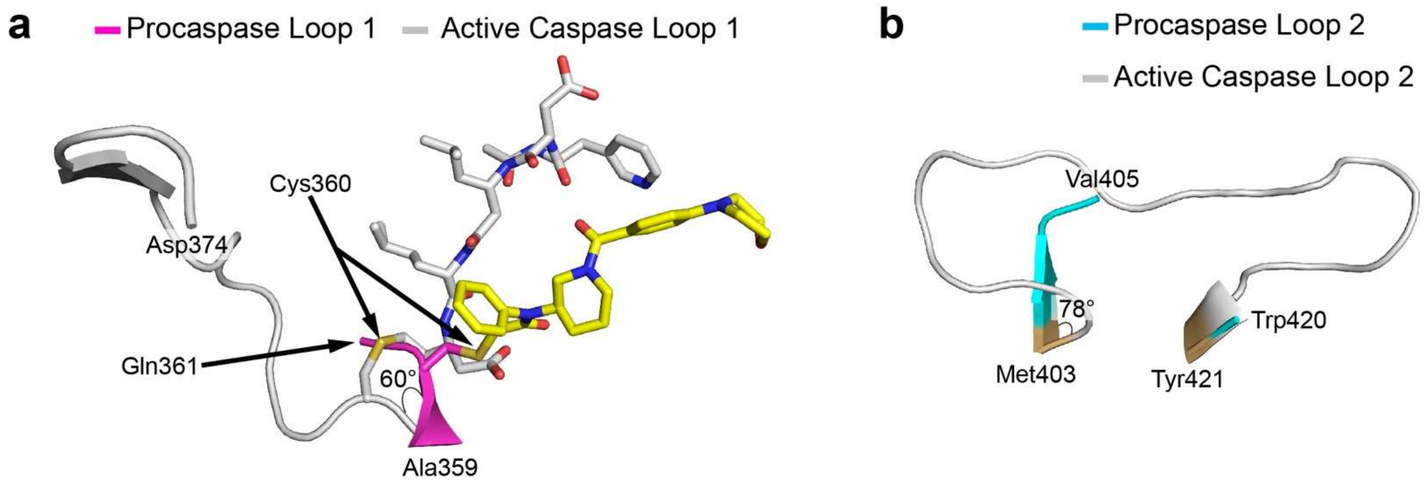


**Supplementary Fig. 1**. Overlay of pro- and active caspase-8 loops 1 and 2. **a** Cartoon representation of the N-terminal end of loop 1 where the catalytic Cys360 of active caspase-8 repositions 60° from Cys360 in the procaspase-8 structure (carbons are magenta, grey, yellow, and green for procaspase-8, active caspase-8, ***63-R***, and active caspase-8 peptide inhibitor respectively, with blue nitrogens, and red oxygens). **b** Cartoon representation of loop 2 where procaspase-8 is cyan and active caspase-8 is grey. Met403 is shifted 78° in the activated caspase-8, changing the secondary structure of loop 2 from the β-sheet seen in procaspase-8 into a disordered loop.

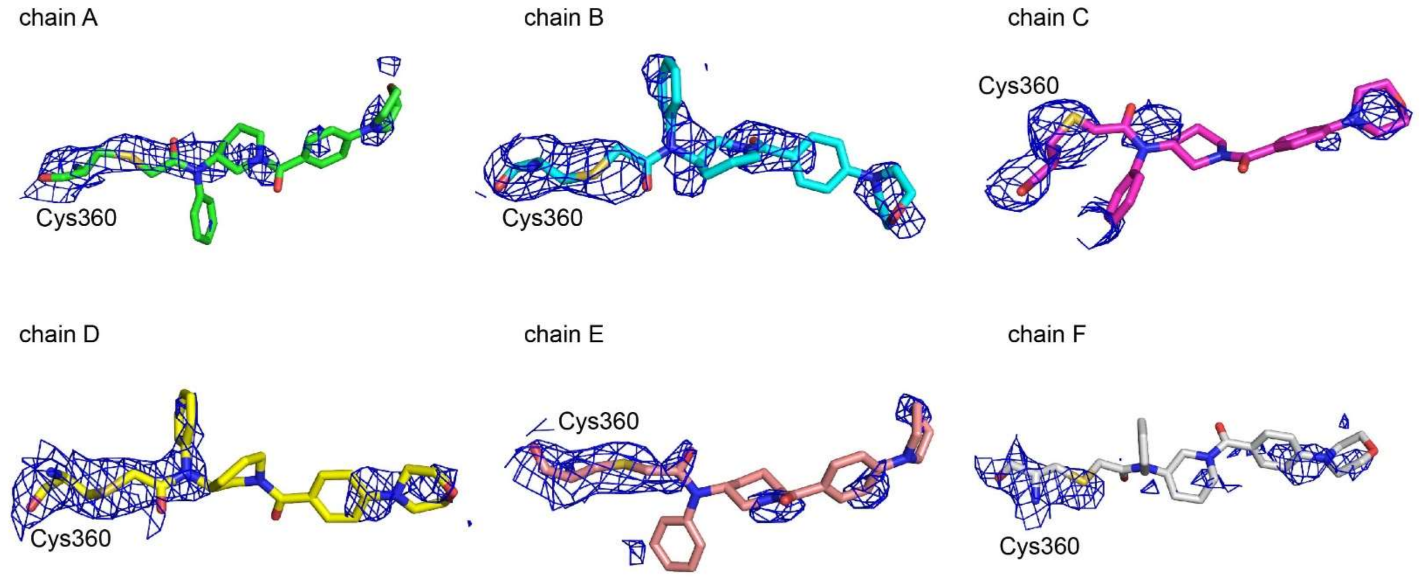


**Supplementary Fig. 2**. Simulated-annealing omit map density contoured at 1.0$\sigma$ of catalytic Cys360 bound to inhibitor **63-*R*** in all 6 subunits. Atoms colored as Supplementary Fig. 1.

**Supplementary Table 2.** Modification of procaspase-8 crystals by **63-*R*.** Crystals from procaspase-8 co-crystallized with **63-*R*** were harvested, reduced, alkylated, subjected to trypsin digest and analyzed by LC-MS/MS. Underline marks the **63-*R***-modified cysteine.

| Protein | Cysteine | Fragment # | Peptide | M+H calculated (m/z) | M+H observed (m/z) | Charge |
|---|---|---|---|---|---|---|
| CASP8 | C360 | 63-*R* | K.VFFIQA<u>C</u>QGDNYQK.G | 1034.99 | 1034.99 | +2 |

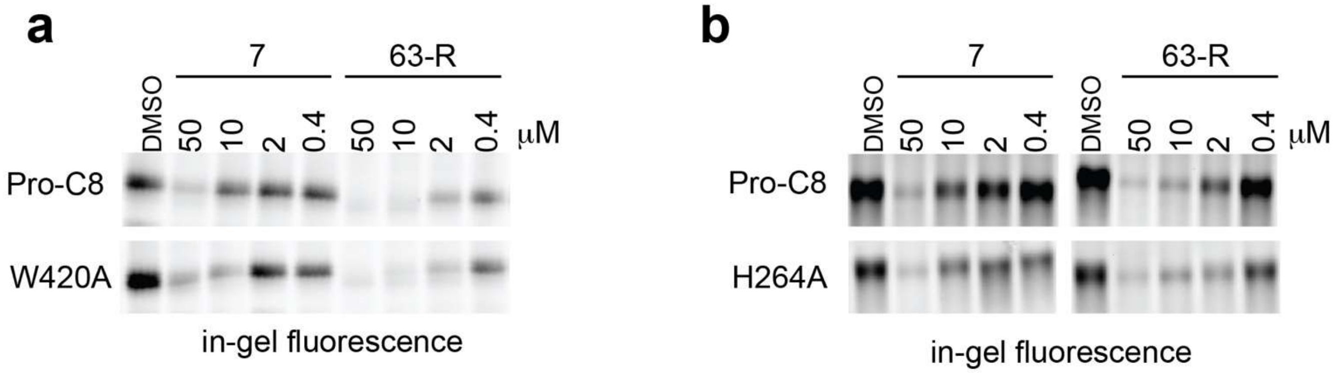


**Supplementary Fig. 3.** Competitive ABPP gels of the W420A and H264A mutated forms of procaspase-8. **a** Recombinant procaspase-8 (D384A and D394A), and mutant procaspase-8 proteins (W420A) were added to HEK 293T soluble lysates to a final protein concentration of 500 nM. The protein-containing lysates were then treated with **7** or **63-*R*** at the indicated concentrations or vehicle (DMSO) for 1h, followed by labeling with **61** (10 μM) for 1h, "click" conjugation to rhodamine-azide, and analysis by SDS-PAGE and in-gel fluorescence. **b** As in '**a**' but with the H264A mutant of procaspase-8. Due to observed instability of the H264A protein upon multiple freeze thaw cycles, the protein was assayed in *E coli* lysates after overexpression without freezing and without further purification.

**a**

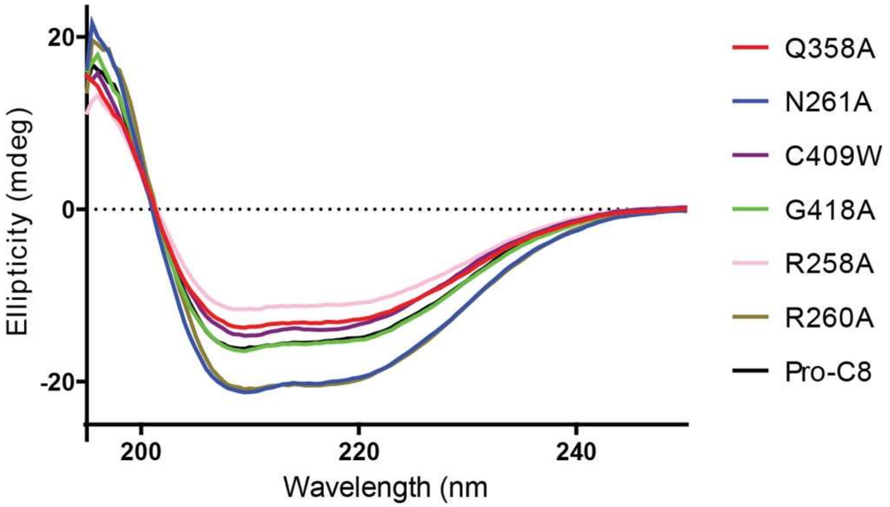


| | Alpha Helix | Beta Sheet | Turn | Other |
|---|---|---|---|---|
| Pro-C8 | 15.9% | 38.3% | 13.9% | 33.4% |
| R260A | 15.3% | 42.5% | 14.9% | 29.3% |
| Q358A | 15.0% | 42.5% | 13.9% | 30.6% |
| G418A | 15.1% | 38.5% | 11.1% | 36.5% |
| C409W | 14.6% | 40.2% | 12.0% | 34.0% |
| R258A | 15.9% | 36.9% | 13.5% | 34.8% |
| N261A | 15.4% | 42.5% | 13.4% | 30.1% |

**b**

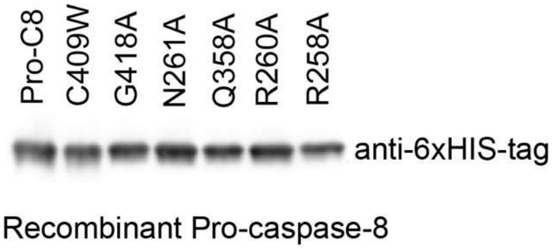


**Supplementary Fig. 4. a** Circular dichroism spectra and calculated secondary structures of caspase-8 mutant proteins. **b** Relative abundance of the indicated recombinant procaspase-8 constructs was visualized by Western blot with an anti-his antibody.

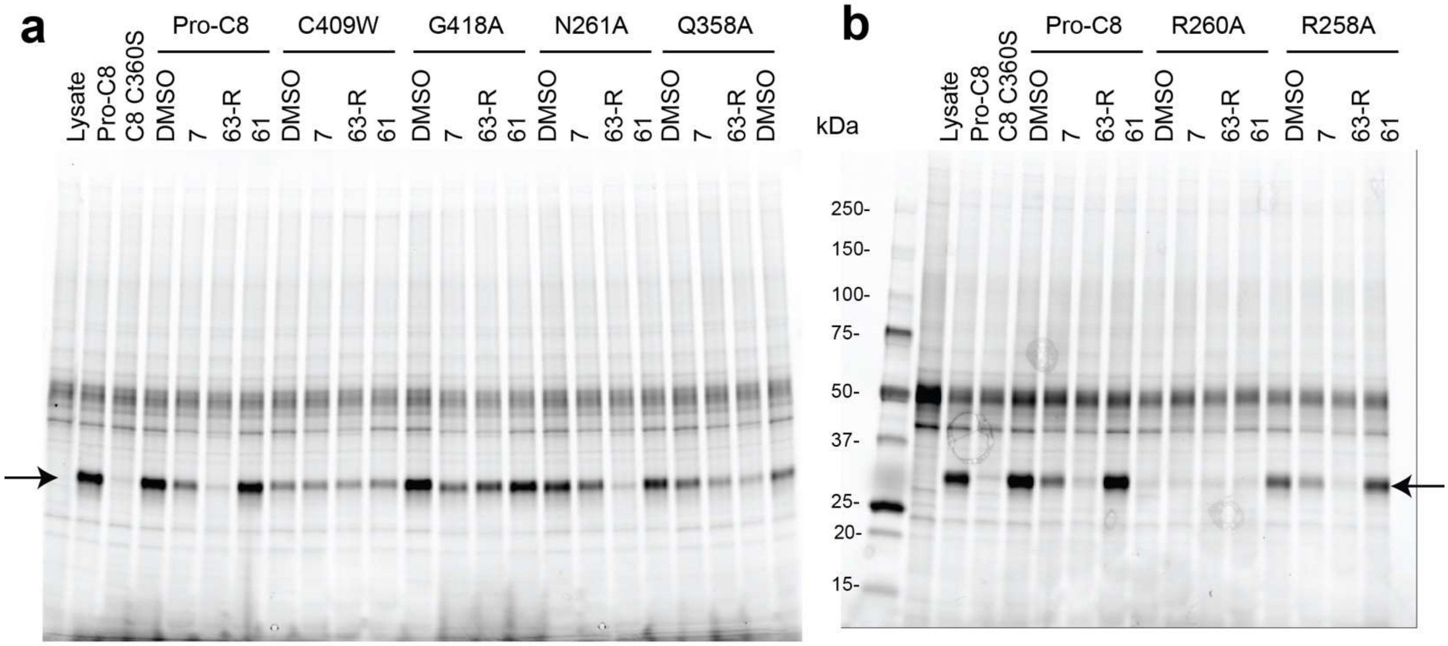


**Supplementary Fig. 5.** Representative full-length gels for single dose competitive labeling experiments quantified in Fig. 5B-I. **a** Recombinant procaspase-8 (D384A and D394A), and mutant procaspase-8 proteins (C409W, G418A, N261A, or Q358A) were added to HEK 293T soluble lysates to a final protein concentration of 500 nM. The protein-containing lysates were then treated with **7**, **63-*R***, **62** (all at 10 μM), or vehicle (DMSO) for 1h, followed by labeling with **61** (10 μM) for 1h, "click" conjugation to rhodamine-azide, and analysis by SDS-PAGE and in-gel fluorescence. Note that the C360S-mutant of procaspase-8, which lacks the catalytic cysteine, did not label with **61**. **b** As in '**a**' but with the R260A- and R258A-mutants of procaspase-8. Arrows indicate procaspase-8 band.

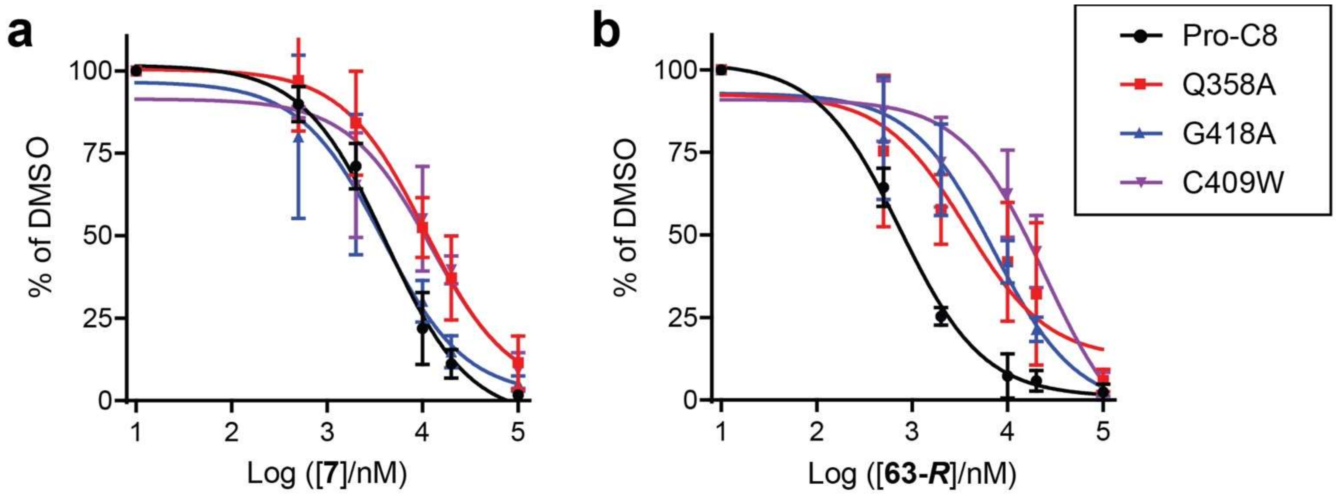


c

| | Pro-C8 | | C409W | | G418A | | Q358A | |
|---|---|---|---|---|---|---|---|---|
| Compound | **7** | **63-*R*** | **7** | **63-*R*** | **7** | **63-*R*** | **7** | **63-*R*** |
| Apparent $IC_{50}$ (μM) | 4.10 | 0.75 | 12.18 | 24.59 | 3.80 | 7.14 | 10.59 | 3.64 |
| 95% CI (μM) | 2.94–5.80 | 0.62–0.94 | 3.37–32.80 | 10.53–63.83 | 1.55–9.13 | 3.12–14.70 | 5.15–20.85 | 0.63–24.98 |

**Supplementary Fig. 6**. Site-directed mutagenesis studies to identify residues that determine compound binding to procaspase-8. (**a** and **b**) Apparent $IC_{50}$ curves for blockade of **61** labeling of procaspase-8 (pro-C8) harboring the indicated mutations by pre-treatment with **7** (**a**) or **63-*R*** (**b**). **c** Calculated apparent $IC_{50}$ values, including 95% confidence intervals derived from the three replicate experiments shown in **a** and **b**.

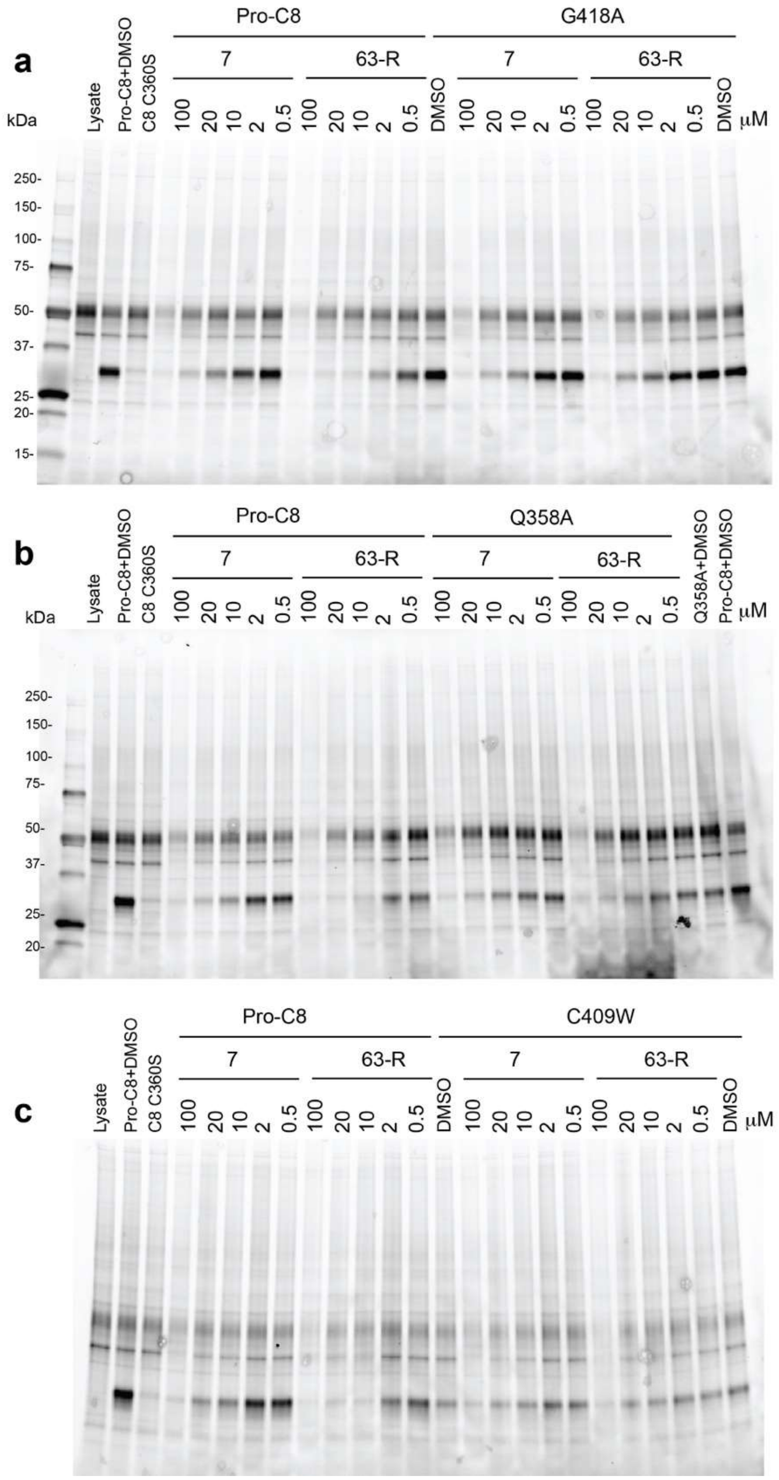


**Supplementary Fig. 7.** Representative full-length gels for $IC_{50}$ competitive labeling experiments quantified in Fig. 5J and Fig. 5 supplement 3. **a** Recombinant procaspase-8 (D384A and D394A), and C409W-mutant procaspase-8 were added to HEK 293T soluble lysates to a final protein concentration of 500 nM. The protein-containing lysates were then treated with **7**, **63-*R***, **62** at the indicated concentrations, or vehicle (DMSO) for 1h, followed by labeling with **61** (10 μM) for 1h, “click” conjugation to rhodamine-azide. **b** as in ‘**a**’, with Q358A-mutant pro-caspase-8. **c** as in ‘**a**’, with C409W-mutant procaspase-8.

### Supplementary References


1. Weiss, M. S. Global indicators of X-ray data quality. *J. Appl. Crystallogr.* **34,** 130–135 (2001).
2. Weiss, M. S. & Hilgenfeld, R. On the use of the merging R factor as a quality indicator for X-ray data. *J. Appl. Crystallogr.* **30,** 203–205 (1997).
3. Karplus, P. A. & Diederichs, K. Assessing and maximizing data quality in macromolecular crystallography. *Curr. Opin. Struct. Biol.* **34,** 60–68 (2015).
